# Age-associated gut microbiota impairs hippocampus-dependent memory in a vagus-dependent manner

**DOI:** 10.1101/2021.01.28.428594

**Authors:** Damien Rei, Soham Saha, Marianne Haddad, Anna Haider Rubio, Blanca Liliana Perlaza, Marie-Noelle Ungeheuer, Harry Sokol, Pierre-Marie Lledo

## Abstract

Aging is known to be associated with hippocampus-dependent memory decline, but the underlying causes of the age-related memory impairment remain yet highly debated. Here we showed that fecal microbiota transplantation (FMT) from aged, but not young, animal donors in young mice is sufficient to trigger profound hippocampal alterations including astrogliosis, decreased adult neurogenesis, decreased novelty-induced neuronal activation and impairment in hippocampus-dependent memory. Furthermore, similar alterations were reported when mice were subjected to an FMT from aged human donors. To decipher the mechanisms involved in mediating these microbiota-induced effects on brain function, we mapped the vagus nerve (VN)-related neuronal activity patterns and report that aged-mice FM transplanted animals showed a reduction in neuronal activity in the ascending VN output brain structure, both in basal condition and following VN stimulation. Targeted pharmacogenetic manipulation of VN-ascending neurons demonstrated that the decrease in vagal activity is detrimental to hippocampal functions. In contrast, increasing vagal ascending activity alleviated the adverse effects of aged mice FMT on hippocampal functions, and had a pro-mnesic effect in aged mice. Thus, pharmacogenetic VN stimulation is a potential therapeutic strategy to lessen microbiota-dependent age-associated impairments in hippocampal functions.

**Graphical abstract:** 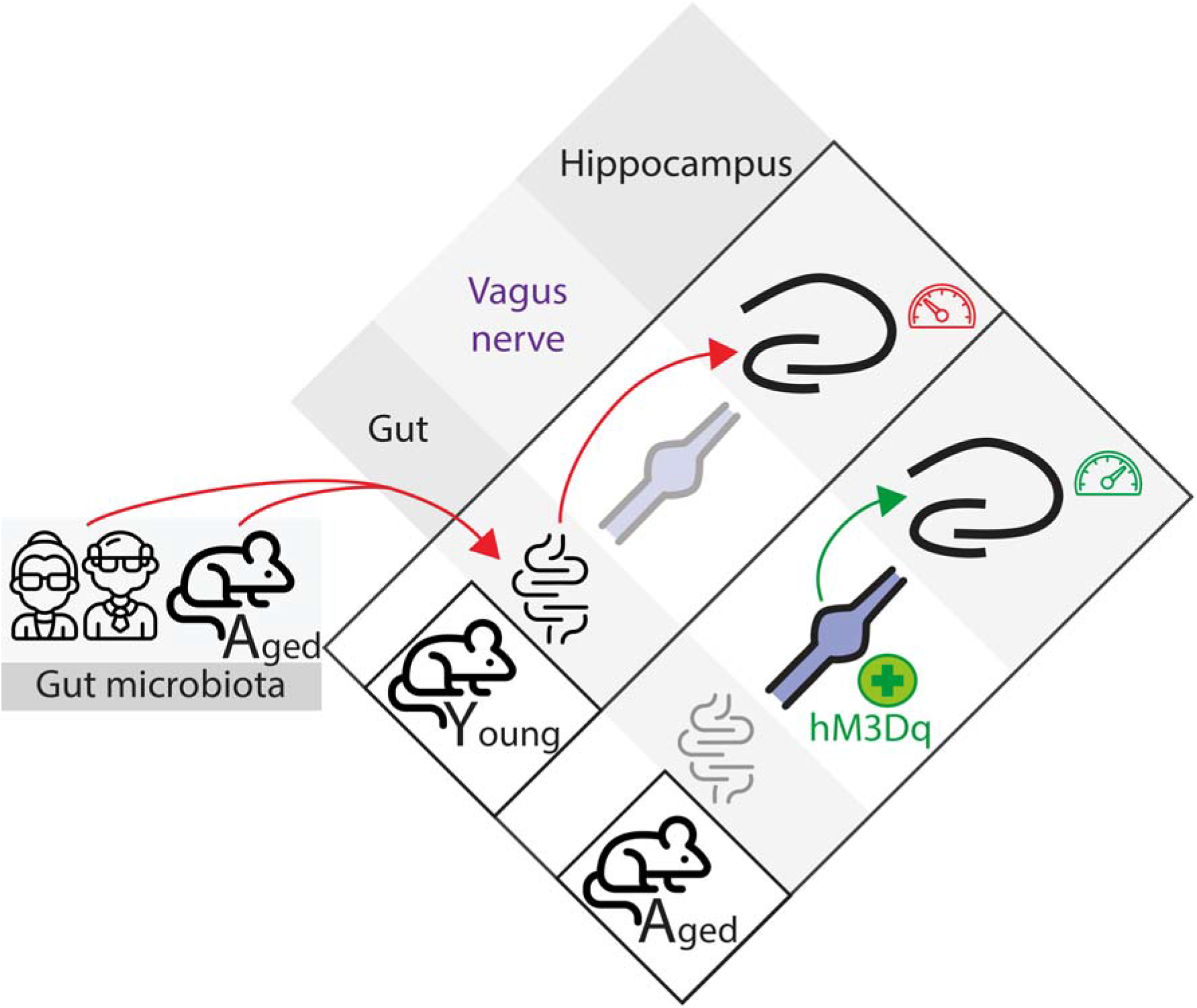

## Introduction

The gut microbiota (GM) – the intestinal community of microorganisms – recently emerged as a key player for physiology and homeostasis, particularly for some brain functions (reviewed in ^1^), notably learning and memory^2–4^. Indeed, the GM influences several hippocampal traits such as adult neurogenesis^4–6^, astrocyte functions^7^ as well as the overall level of brain inflammation^8,2^. Intriguingly, these same brain processes are also affected by aging, altering brain functions and ultimately leading to hippocampal-dependent impairment in episodic and spatial memory^9^. However, understanding the GM’s role on the detrimental effects of the aging process is a daunting task since age influences the microbiota composition. So, it remains unknown to what extent gut microbes contribute to the aging phenotype, and if they do, what are the mechanisms at play.

Evidence is starting to uncover the causal role of GM changes on cognitive functions. In recent studies, GM from aged rodents, transplanted to their young counterparts, impaired learning and memory performances in the Barnes maze^3^ or delayed matching to position tests^2^. It also promotes systemic^8^ and hippocampal inflammation and is associated with a perturbation in the expression levels of synaptic plasticity genes^2^.

Beyond the possible link via inflammation, the GM is also known to directly communicate with the brain, notably through the vagus nerve (VN). The VN circuit is the most direct and well-studied neuronal pathway of the gut-brain axis (reviewed in^10^). VN sensory fibers innervate the muscular and mucosa layers of the gastrointestinal tract, detect mechanosensory and chemical signals, and then relay these signals to the CNS through the ventro-medial part of the nucleus tractus solitarius (vmNTS) located in the caudal brainstem^11^. Interestingly, activity in this vagal gut-brain circuit is known to modulate hippocampus (HPC) function. Indeed, the antidepressant effect of probiotic treatment on the hippocampal expression levels of GABA receptors was shown to require an intact VN, since this effect disappears in vagotomized animals^12^. Furthermore, VN electrical stimulation is an approved therapy for depression and drug-resistant temporal epilepsy (reviewed in ^13^), and is a powerful modulator of hippocampal functions, notably at the electrophysiological^14^ and epigenetic^15^ levels.

Anatomically, the NTS is connected with the HPC through multi-synapses connections, notably through the locus coeruleus and dorsal raphe, which respectively, release noradrenaline (NA) and serotonin (5-HT) to the HPC, and the whole forebrain^16,17^. In line with this, a link between VN ascending neurons and HPC-dependent memory was recently reported using a selective vagal ascending neurons ablation^18^, or Ghrelin receptor knock-down^19^. Both studies report impaired HPC-dependent episodic and spatial memory. However, beyond rodent GM data, the effect of human age-associated GM on hippocampal memory is currently unknown. Here, we report that the transplantation of young mice with an age-associated GM, of both mouse and human origin, transfers some aspects of the deleterious impact of aging on the hippocampus, highlighted by a deficit in memory and an inability of the hippocampal network to respond to a novel environment. Furthermore, targeted pharmacogenetic manipulation of VN activity uncovers the role of VN tone on hippocampal function and its potential therapeutical impact on age-related hippocampal dysfunction.

## Results

### Age-associated GM, of both murine and human origin, impairs hippocampal functions

To investigate the potential contribution of the transplanted GM in the detrimental impact of aging on hippocampal functions, young (**Y**) adult mice (2-month-old, hereafter referred to as “young” mice) were treated with broad-spectrum antibiotics, followed by FMT. This method allowed for the transplantation of the recipient mice with the GM from young (**Y**oung mice GM in **Y**oung mice = **YinY**) or aged (**A**)-(18-month-old) donor mice (**A**ged mice GM in young mice = **AinY**) (Fig. 1A).

**Fig. 1.**
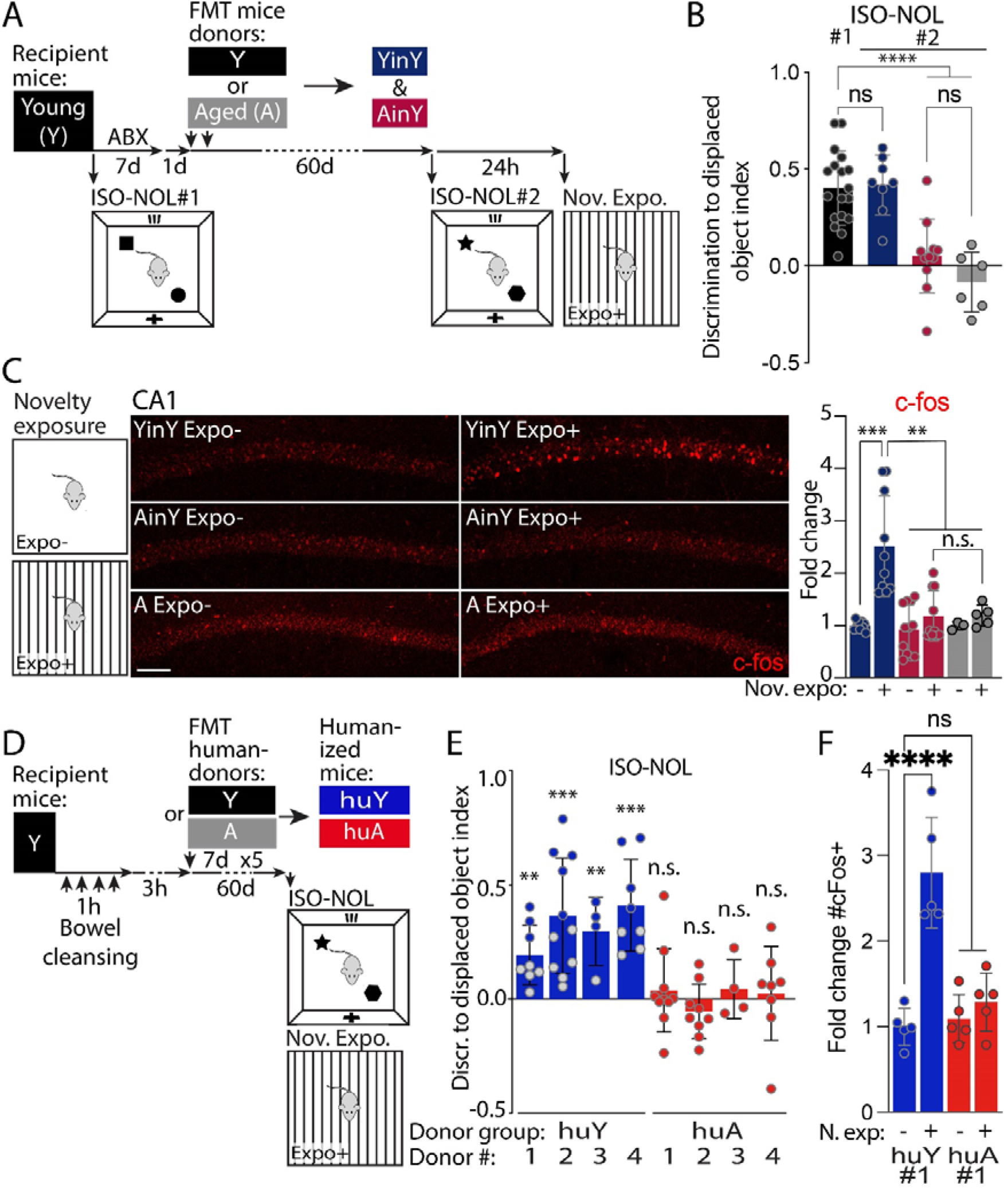
Age-associated GM, of both murine and human origin, impairs hippocampal function and structure. (A) Schematic of the age-associated mouse GM transplantation scheme and behavior in young (Y) adult (2-3 months) mice and, (B) of its effect on memory abilities in the isotropic version of the novel object location task (ISO-NOL) of young mice prior to, and after, GM transplantation from either young (Young mice GM in Young mice = YinY blue bars) or aged (>18 months) animals (Aged mice GM in young mice = AinY) compared to aged mice (n = 18, 8, 10 and 6 mice per group). (C) Representative immunohistochemical images and quantitative analysis of the effect of age, versus young, -associated GM transplantation compared to aged mice on the number of c-fos-positive cells measured in CA1, after exposure to novelty (n = 7, 8, 10, 7, 3 and 5 mice per group). (D) Schematic of the human GM-transplantation scheme and behavior in young mice. (E) Effect of the colonization of young mice with the GM from young human (huY) *versus* aged (HuA) donors on memory abilities in the ISO-NOL behavioral task (n = 4 donors per age human-GM group, with 5 to 10 replicate mice per donor). (F) Quantification of the effect of huA donor#1 *versus* huY donor#1 GM transplantation on recipient-mice’s increase in the number of dorsal hippocampal CA1 c-fos-positive cells after exposure to novelty (n = 5 per group). (B, C and E, F) The color codes of the different quantification experimental groups are depicted in the (A) and (D) schematics. (B, C and F) one-way ANOVA and (E) one-tailed t test. Throughout the figure, bars represent the mean ± SD. N.s., non-significant; p >0.05; ** p ≤ 0.01; *** p < 0.001. Scale bars, 100 μm.

To verify the effectiveness of the antibiotic treatment to cleanse the mice GM prior to FMT, we first analyzed the decrease in bacterial content in a FM plating experiment made from samples collected from the same mice before and after the antibiotic treatment (Fig. S1-1). Then, analysis of the microbial composition from the donors and FM transplanted mice was performed by 16S rRNA gene sequencing. Changes in the relative abundance of bacteria in both donor and recipient mice were observed at the taxonomic level of family (Fig. S1-2A), with a trend toward an age-associated increase in the Firmicutes to Bacteroides ratio in donor samples that was no longer seen post-FMT (Fig. S1-2B). Alpha diversity can be used to measure the richness and uniformity of species in community ecology. Using Shannon and Simpson indexes or Abundance-based Coverage Estimator (ACE), we did not observe significant changes in Alpha diversity between Y, A, YinY and AinY groups (Fig. S1-2C-E). Determination of the beta diversity, the comparative analysis of the composition of the microbial community in different samples, showed differences in the GM from young and old donors (clusters Y and A, Fig. S1-2F) and recipient young mice receiving young- or age-associated microbiota (clusters YinY and AinY, Fig. S1-2F). Although such changes were relatively subtle compared to previous studies using mice from different strains^20–23^, comparative analysis of the young *versus* aged mice and transplanted animals showed a clustering due to the age of the donor that was transferrable to the colonized animals (Fig. S1-2F). This result is supported by the lower Jaccard distance between donor and recipient mice than between Y and A or YinY and AinY mice (Fig. S1-2G). Thus, we conclude that, even though the FMT approach used here led to a distinct microbial composition signature in each of the four different groups, yet some of the GM “aging” characteristics are transferrable to the recipient mice.

Next, the repercussion of GM colonization on hippocampal functions was investigated at the behavioral and cellular levels. Hippocampus-dependent memory abilities was scored before and after GM-colonization using the hippocampus-dependent isotropic version of the novel object location (ISO-NOL) task^24,25^, a task sensitive to aging^26^. In this assay, young mice showed normal discrimination memory performances in the test before FMT, and no changes were observed following GM-transplantation from young donors (Fig. 1B). Conversely, GM-transplantation from aged mice led to a deterioration of the recipient mice’s memory to levels similar to the scores seen in naïve-aged mice (Fig. 1B). The detrimental impact of the aged mice-GM on young recipient mice memory was not specific to this task, as similar deficits were also seen in other hippocampus-dependent behaviors, *i.e*., the novel object recognition^27^, in a version where the dorsal HPC implication was previously demonstrated in this task^28^, and the contextual version of the fear conditioning^29^, to a level of severity that matched the deficits seen in naïve-aged animals (Fig. S1-3A-C). As the behavioral tasks use here rely on different HPC functions/connectivity networks: *i.e*., the HPC solely for the ISO-NOL task, and HPC connection to the perirhinal and insular cortexes and to the amygdala for the ISO-NOR^27^ and fear conditioning^30^, respectively, it seems that the age-associate GM negatively impact the ensemble of HPC memory.

To examine the impact of the age-associated GM on the hippocampus neuronal network, alterations in the expression of the immediate early gene c-fos following novelty exposure^31,32^ were quantified by automated counting in the dorsal hippocampal CA1 (Fig. 1C). This subregion was selected due to its known involvement in the memory task used here, an implication demonstrated through inactivation experiments or electrophysiological recordings^24,33,34^. Novelty-induced upregulation of c-fos in the dorsal CA1 was seen in animals colonized by a young GM, while mice receiving an age-associated microbiota showed reduced activation similar to that of aged mice (Fig. 1C). A comparable deficit in the density of c-fos-positive neurons in response to mice exposition to novelty was also observed in the CA3 and dentate gyrus (DG) HPC subregions of age-associated GM transplanted mice (Fig. S1-3D).

Then, we analyzed hippocampal astrogliosis and adult neurogenesis, two cellular processes highly sensitive to aging^35^ and to GM composition^36^. As depicted in Fig. S1-3E, age-associated GM transplantation increases astrogliosis. It also decreases significantly adult hippocampal neurogenesis as reported by a reduction in the number of DCX-expressing immature adult-born neurons in the DG (Fig. S1-3F). Thus, age-associated GM transplantation to young animals impairs hippocampal memory, the ability of the HPC network to respond to novelty exposure, promotes astrogliosis and decreases the number of hippocampal newly-generated neurons, mimicking important hallmarks of aging on hippocampal structure and function.

We then sought to investigate whether the transplantation of human gut microbiota could promote similar changes in the hippocampus of recipient mice. Fecal samples were collected from adult healthy human donors classified as young (less than 35 years old), or aged (more than 65 years old). Following human-FMT into young recipient mice, the potential effects on memory were evaluated using the ISO-NOL task (Fig. 1D). Mice transplanted with young human-GM perform normally in the task. In contrast, the microbiota transplantation from aged human donors impaired hippocampus-dependent memory (Fig. 1E), and prevented the novelty-induced CA1 activation (Fig. 1F and S1-3G). Therefore, the microbiota transplantation from old, but not young, human donors promotes alterations in hippocampal function in the recipient mice. Together, these data show that the age-associated GM, of both murine and human origin, plays an important role in the etiology of age-associated memory deficits.

### Decrease in vagus nerve activity is both necessary and sufficient for age-associated GM impact on the hippocampus

To investigate whether the HPC deleterious impact of age-associated mouse GM transplantation in young mice depends on VN signaling, the c-fos-based activity level was assessed in the vmNTS, the entry point of VN ascending signaling into the brain (schematic Fig. 2A), of age-associated GM transplanted young mice. The number of c-fos-positive neurons in the vmNTS was taken as a proxy of VN ascending activity level. Automated quantification of c-fos-positive cells in this brain region revealed a reduction in neuronal activity in age-associated GM transplanted mice compared to their young-associated GM transplanted counterparts. This holds true under basal conditions (Fig. 2B) and following VN activation through food and water consumption (Fig. 2C). Such analysis reveals that age-associated GM transplantation leads to a deficit in VN ascending inputs to the brain, both at rest and following VN activation. Next, to directly test the possible implication of VN activity for hippocampal memory function, we manipulated VN ascending activity by viral transduction of the designer receptors exclusively activated by a designer drug (DREADD) in the left nodose ganglia (L-NG). This side was chosen to be consistent with the side used for electrical VN stimulation as a treatment for resistant major depression^37^. Co-injection of AAV1-hSyn-Cre and AAV5-hSyn-DIO-hM3Dq-mCherry efficiently labelled L-NG neurons soma and their axonal terminals could be detected in the ventromedial areas of the NTS (Fig. 3A). Three weeks post-viral delivery, the DREADD activator clozapine N-oxide (CNO) was administered (Fig. S2A). Using the excitatory form of DREADD, we confirmed that CNO injection significantly increased the number of c-fos-positive neurons in the vmNTS (Fig. S2B-C). Furthermore, when co-expressing both excitatory and inhibitory DREADD, we confirmed that the inhibitory DREADD hM4Di in the L-NG blocks such CNO-induced increase in vmNTS c-fos expression (Fig. S2B-C). This result demonstrates that our experimental procedure enables both activatory and inhibitory pharmacogenetic-control of VN ascending signaling.

**Fig. 2.**
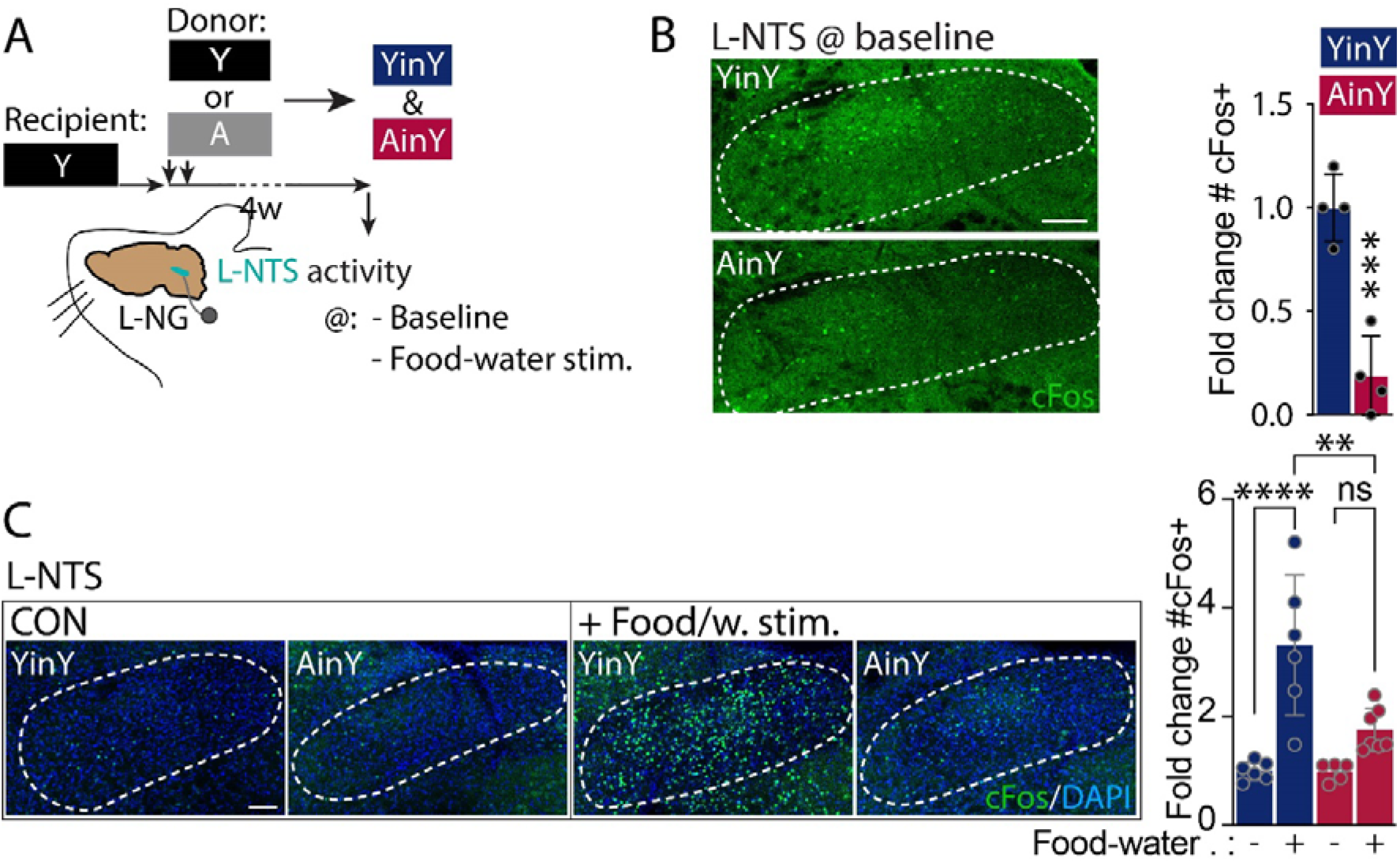
Impact of age-associated GM transplantation on ascending VN signaling. (A) Schematic of the left nodose ganglia (L-NG) projections to the left nucleus of the solitary tract (L-NTS), and effect of the age-associated GM transplantation on the number of c-fos positive cells in the NTS at baseline condition (B, n = 4 per group) and following food and water consumption (C, n = 6, 6, 5 and 7 per group). (B) Student’s t test, (C) one-way ANOVA. Throughout the figure, bars represent the mean ± SD. N.s., non-significant; p > 0.05; * * p ≤ 0.01; * * * p > 0.001, * * * * p > 0.0001. Scale bars, 100 μm.

**Fig. 3.**
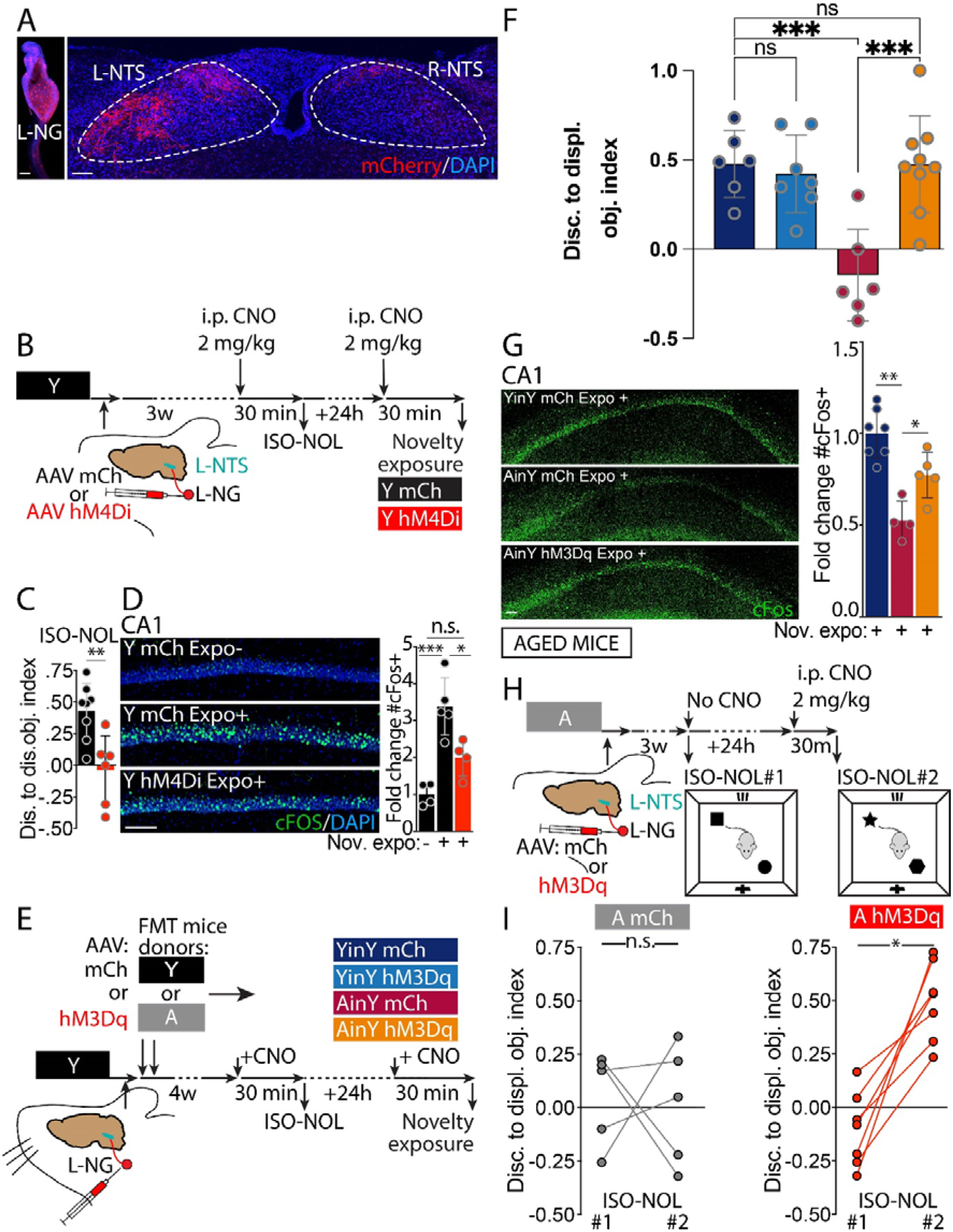
Decrease in VN signaling is necessary and sufficient for the age-associated GM negative impact on memory and activation of VN ascending signaling increases the memory abilities in aged mice. (A) Representative image of an AAV-mCherry transduced left nodose ganglia and of its ascending mCherry-positive fibers in the left (L)-and right (R) -NTS. (B, E and H) Schematic of the viral and pharmacogenetic-based strategy to modulate VN ascending signaling in young (B, E) and aged (H) mice where mice L-NG were AAV1-hSyn-Cre (not represented) and hM4Di -mCherry (B) or hM3Dq-mCherry (E, H), or AAV5-hSyn-DIO-mCherry for control mice, co-injected, and timing for the behavior and pharmacogenetic ascending VN activity modulations. Effect of L-NG DREADD inhibition on, (C) the memory ability in the ISO-NOL task (n = 8 and 6) in L-NG hM4Di inhibited versus control mCherry-only mice and, (D) the increase in the number of dorsal hippocampal CAI c-fos positive cells after exposure to novelty in L-NG hM4Di inhibited versus mCherry-only control mice (n = 4, 5 and 4 per group). Effect of the pharmacogenetic vagal activation in AinY mice and its effect on, (F) the memory abilities in the novel object location task (n = 6, 7, 5 and 8) and, (G) the increase in the number of dorsal hippocampal CAI c-fos-positive cells after exposure to novelty, in YinY and AinY mCherry-only versus AinY hM3Dq expressing mice (n = 7,4 and 5 per group). (I) Effect of L-NG DREADD activation on ISO-NOL memory ability in L-NG mCherry or hM3Dq expressing aged mice, without (ISO-NOL#l) versus with (ISO-NOL#2) ascending VN activation (n = 5 and 7). The color codes of the different experimental groups are depicted in the associated schematics in (B), (E) and (H). (C) Student’s t test, (D, F and G) one-way ANOVA, (I) two-tailed t test with Wilcoxon matched-pairs signed rank test. Throughout the figure, bars represent the mean ± SD. N.s., non-significant; p > 0.05; * p < 0.05; * * p > 0.01; * * * p > 0.001. Scale bars, 100 μm.

We next evaluated the hippocampal memory strength following a transient inhibition of the vagal ascending activity, thus mimicking the age-associated GM effect on ascending vagal signaling. L-NG inhibition during the ISO-NOL test (Fig. 3B) completely impaired mice’s discrimination ability in the task (Fig. 3C) and this loss was associated with an absence of novelty-induced increase in c-fos-positive cell number counted in the dorsal CA1 region (Fig. 3D). We next investigated whether a stimulation of ascending VN activity could counteract the age-associated GM deleterious effect. To test this possibility, L-NG of YinY and AinY mice were transduced to express the excitatory hM3Dq DREADD (Fig. 3E). Pharmacogenetic VN activation (Fig. S2D), resulted in the induction of c-fos expression in the L-NTS of the hM3Dq-transduced GM groups in comparison to mCherry-only control animals (Fig. S2E), demonstrating the robustness of the protocol used to activate the ascending VN pathway, irrespectively of the transplanted-GM donor’s age. In the ISO-NOL memory task, we showed that hM3Dq L-NG transduced YinY mice were not affected by the manipulation, while hM3Dq-AinY mice depicted a complete rescue of their memory abilities, when compared to AinY and YinY, mCherry controls (Fig. 3F). Thus, pharmacogenetic VN stimulation can reinstate normal cognitive status in AinY mice. At a cellular level, this DREADD ascending-VN stimulation increases hippocampal c-fos induction following animal exposure to novelty in L-NG hM3Dq transduced AinY mice when compared to AinY and YinY mCherry-only controls (Fig. 3G). The latter result **shows that the increase in ascending VN signaling rescues the HPC ability to respond to novelty** in AinY mice. Collectively, these data reveal that **inhibiting the activity of ascending VN activity is necessary and sufficient** for the detrimental impact of age-associated GM transplantation on HPC memory.

Finally, we reasoned that since age-associated GM inhibition of VN ascending signaling was causal to the detrimental impact of aging on memory, thus restauration of its activity should also restore memory of aged mice. This hypothesis was tested by performing ascending-pharmacogenetic VN activation in aged mice (Fig. 2H). This manipulation indeed increased the memory ability of CNO-activated hM3Dq transduced aged mice in the ISO-NOL test, in comparison to levels of the same animal at baseline (prior day with no CNO injection), whereas no memory enhancement was seen in control mCherry-only VN transduced aged animals (Fig. 2I). **These findings open new therapeutic approaches aimed at acting either, directly on the GM composition, or on VN activity, to alleviate the detrimental impact of aging on memory**.

## Discussion

In conclusion, our study unravels a contribution of age-associated GM —from both murine and human origins— in mouse hippocampal alterations similar to those observed with aging. It also highlights the participation of the VN in mediating this age-related phenomenon through the gut-brain axis (Fig. S3). To translate the effect of an acute pharmacogenetic modulation of NG ascending activity to the HPC, a neural transduction pathway is likely to take place. Indeed, given the temporal dynamic, the neural route seems more probable, as opposed to the humoral and/or immunity-based routes of communication. It is particularly interesting to note that this acute pharmacogenetic ascending VN activation can reinstate normal memory in age-associated GM transplanted and aged mice, despite the presence of GFAP-based signs of HPC neuroinflammation in these mice. We postulate that an increase in VN ascending signaling might counteract the neuroinflammation to reinstate a normal hippocampal function. This neural pathway hypothesis is consistent with the fact that the VN connects both the locus coeruleus and dorsal raphe, which release noradrenaline (NA) and serotonin (5-HT), respectively^16,17^. The VN also exerts a modulation in activity of the basal forebrain area^38^, which releases acetylcholine (Ach) to the entire forebrain, including the HPC. Therefore, a deficit in ascending NG could lead to a reduction of those neurotransmitters release in the HPC. This would lead to a deficit in the general activity level in the HPC, that would translate into a deficit in memory abilities. This hypothesis supports the decrease in neuronal activity level seen in the HPC of age-associated GM microbiota transplanted animals in the novelty exposure assay used here. This NTS/basal forebrain modulation of activity was recently illustrated with the description of the NTS-Septum-HPC circuit regulation of hippocampal-dependent memory^18^. The fact that a stimulation of VN ascending signaling is effective in counteracting the detrimental impact of both: 1) the age-associated GM transplantation and, 2) aging, raises the question of the specificity of this procedure. This is an interesting question that is beyond the scope of our study but should be addressed in future research. For instance, this question could be tackled by testing the effect of the pharmacogenetic-mediated increase in VN ascending signaling in some microbiota-related contexts distinct from aging, like in a model of antibiotic driven hippocampal memory dysfunction^4^, or microbiota independent paradigms such as acute pharmacologic retrograde amnesia^39^. As for the potential neuronal circuit involved in translating the age-associated GM effect through the vagal/NTS system to the HPC described herein, this communication route must be multi-synaptic. Indeed, the vmNTS, where the VN projects its terminals, connects to many brainstem and forebrain regions, but not to the HPC directly^40,41^. Beside the possible involvement of the NTS-septum-HPC circuit mentioned above^18^, another possibility is the recently described connection of the locus coeruleus with the HPC through dopaminergic connections^42^. This brain structure is known to be one of the main relay of vagal activity and its dopaminergic projections to the HPC were recently described to be prominent in the neural encoding of novelty by the HPC^42^. This function and circuit are most likely important in the detection of novelty in the hippocampal memory tasks used here, such as the NOL-memory test, and the possible involvement of a decrease in LC-dopaminergic inputs to the HPC through the age-associated GM downregulation of VN ascending signaling could be investigated in the future. Notwithstanding the questions that remain unsolved, our findings open innovative therapeutic avenues aimed at acting either, directly on the GM composition, or on VN activity, to alleviate the detrimental impact of aging on memory.

## Methods

### Study approval

<u>Humans:</u> FM (fecal matter) donors that matched the young and age category were selected from the Institut Pasteur (IP) ICAReB plateform’s healthy volunteer Diagmicoll and CoSImmGEn cohorts. The clinical research protocols for collecting and handling of biological material and associated clinical and biological information were reviewed and approved by a French ethical committee “comité de protection des personnes lle de France 1” (Ref: 09-12179, Ref: 2010-dec 12483) and is compliant with European general data protection regulations (GDPR). Donors received an oral and written information about the research and written informed consent was obtained from all recruited human subjects. <u>Animals</u>: experiments were performed using adult (> 10-week-old) male RjOrl: SWISS mice purchased from Janvier labs (St Berthevin, France). Animals were housed under a 12h/12h light/dark cycle, with dry food pellets and water accessible *ad libitum*, at the Pasteur Institute animal care facility, officially registered for experimental studies on rodents. All animal experiments were designed according to the European community council directive of 24 November 1986 (86/609/EEC) and the European Union guidelines, to the 3R’s rules and were supervised by the French Ministry of Research, as well as reviewed and approved by the Animal Welfare Committee of the IP (project number: 2016-0023) and the “Service prevention des risques” (experimental protocol 17.029).

## Author contributions

DR designed, developed, and managed the project, performed all experiments and wrote the manuscript, under the supervision of PML. HS performed 16S rRNA gene sequencing analysis with the help of SS. MH and AHR were involved in behavioral and histological analysis. The ICAReB platform handled the donor samples regulatory and logistic aspects of the project under MNU supervision.

## Acknowledgments

The authors thank the study participants, physicians (Joёlle Brachat, Joёl Ankri, Bertrand Denis, Dany Vythilingum), clinical/research assistants (Caroline Roussel, Céline Nemeth) and health officers (Anaïs Perilhou) who helped conduct this study, Laurence Motreff and Marc Monod from the Biomics Platform, C2RT, IP, Paris, France, supported by France Génomique (ANR-10-INBS-09-09), the life insurance company “MTRL” and IBISA; Blanca Liliana Perlaza, Amina Ait Saadi, Linda Sangari, Sophie Chaouche, Remy Artus from ICAReB; Oriana Lavielle, Gabriel Lepousez, Kurt Sailor and Ilana Gabanyi for reading the manuscript.

