## Supplementary figures and methods for "Age-associated gut microbiota impairs hippocampus-dependent memory in a vagus-dependent manner"

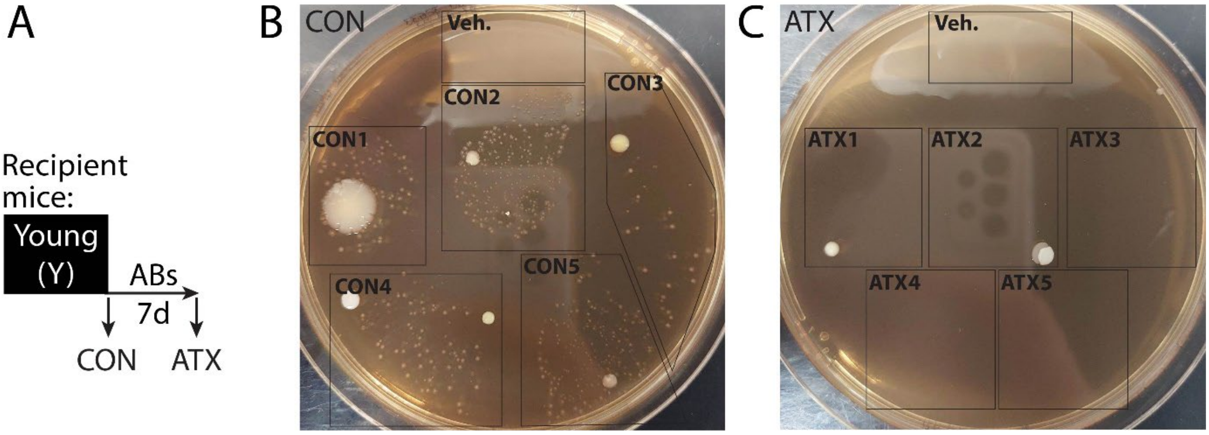


**Fig. S1-1. Antibiotic-based GM cleansing prior to FMT.** (A) Schematic of the experimental procedure to assess the efficiency of the pre FMT large spectrum antibiotic (ABX)-based GM cleansing procedure. Bacterial growth from FM resuspension from the same five mice before (B), and after (C), receiving the antibiotic treatment.

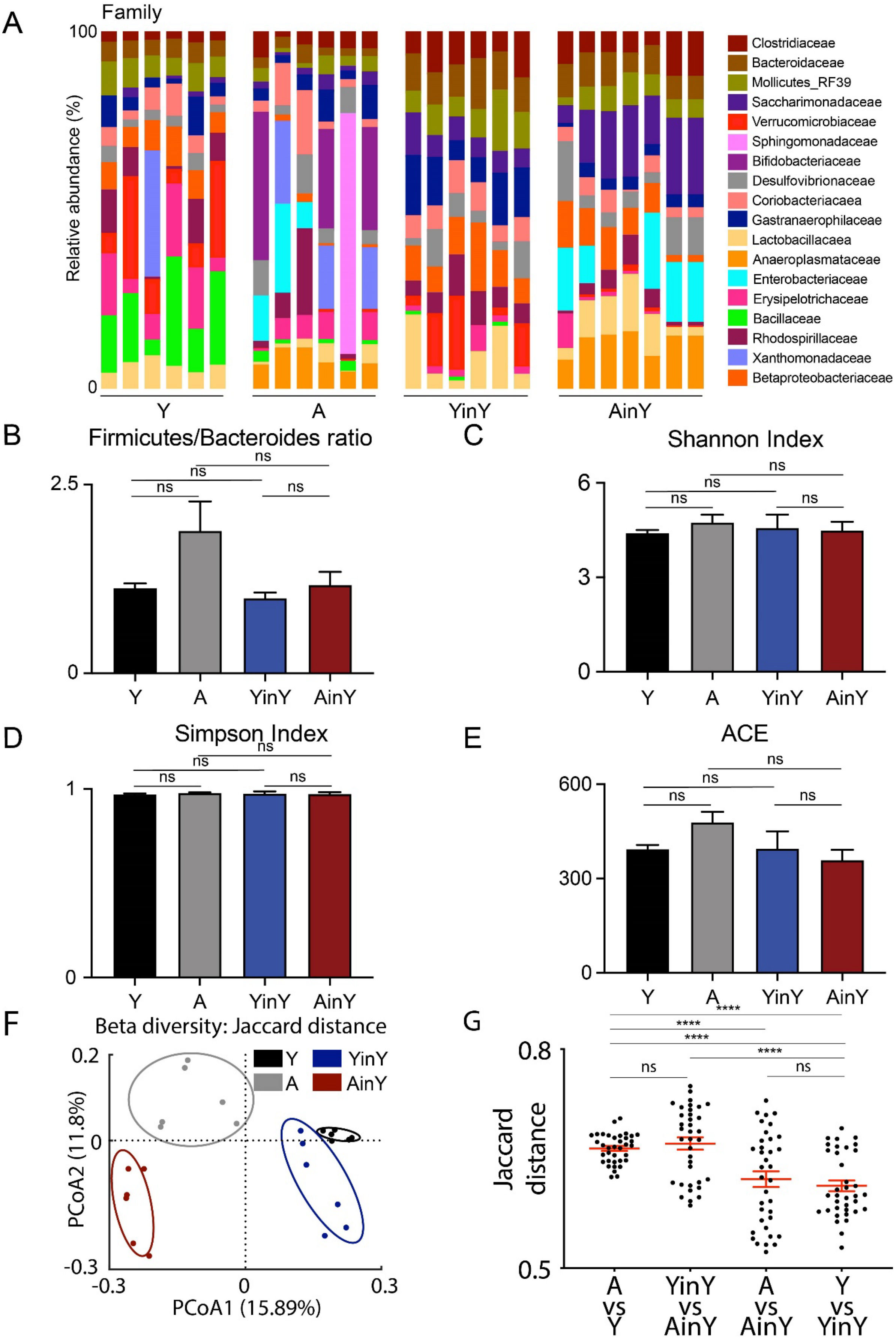

A
Family
100
0
Relative abundance (%)
Y
A
YinY
AinY
Clostridiaceae
Bacteroidaceae
Mollicutes_RF39
Saccharimonadaceae
Verrucomicrobiaceae
Sphingomonadaceae
Bifidobacteriaceae
Desulfovibrionaceae
Coriobacteriacaea
Gastranaerophilaceae
Lactobacillacaea
Anaeroplasmataceae
Enterobacteriaceae
Erysipelotrichaceae
Bacillaceae
Rhodospirillaceae
Xanthomonadaceae
Betaproteobacteriaceae
B
Firmicutes/Bacteroides ratio
2.5
0
ns
Y
A
YinY
AinY
C
Shannon Index
6
3
0
ns
D
Simpson Index
1
0
ns
E
ACE
600
300
0
ns
F
Beta diversity: Jaccard distance
0.2
0
-0.3
PCoA2 (11.8%)
-0.3
0
0.3
PCoA1 (15.89%)
Y
A
YinY
AinY
G
****
ns
0.8
0.5
Jaccard distance
A vs Y
YinY vs AinY
A vs AinY
Y vs YinY

**Fig. S1-2. 16S-based characterization of the age-associated mouse GM-transplantation to young mice model.** (A) Bacterial microbiota composition at the family level in experimental conditions of young and aged donors and transplanted mice (Y, A, YinY and AinY). (B) The Firmicutes-Bacteroides ratio between the experimental groups show no significant changes. Shannon (C) and Simpson (D) index of species diversity, and abundance-based Coverage Estimator (ACE) (E), showed non-significant differences between the four experimental groups. (F) Principal coordinate analysis of Jaccard distance among samples from the four experimental groups, where PCoA1 and PCoA2 represented the top two principal coordinates that captured most of the diversity. The fraction of diversity captured by each coordinate is given as a percentage. (G) Jaccard distance between samples of indicated experimental groups. (B-E, G) One-way ANOVA. Throughout the figure, bars represent the mean ± SD. N.s., non-significant and ****$p < 0.0001$.

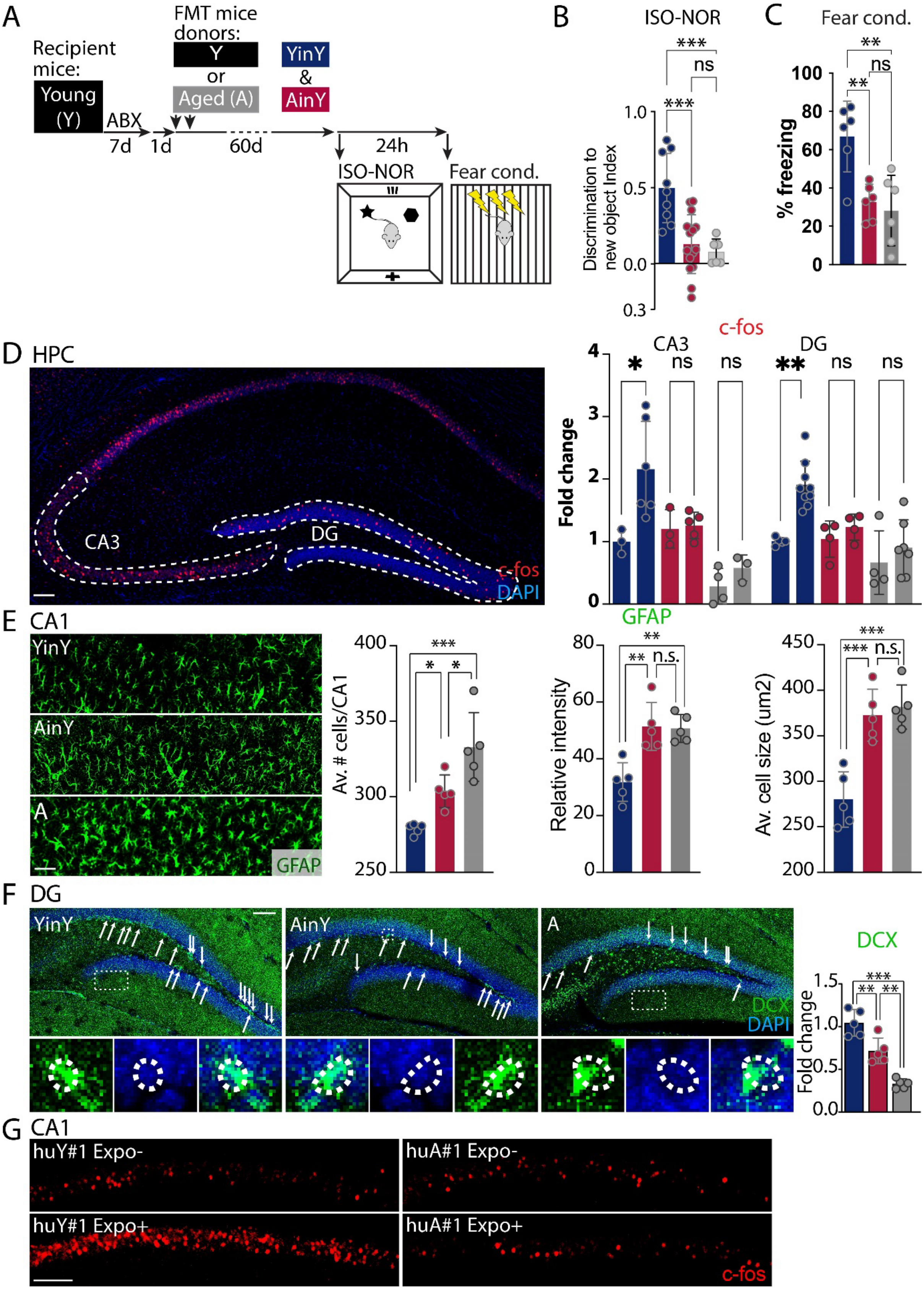


**Fig. S1-3. Age-associated GM impairs hippocampal function in the contextual fear conditioning and novel object-recognition (NOR) tasks.** (A) Schematic of the age-associated GM transplantation scheme in young mice and of its effect on memory abilities of the transplanted mice in, (B) the isotropic version of the NOR task (ISO-NOR; n = 9 and 15 mice per group), and (C) contextual fear conditioning (fear cond., n = 6 per group). (D)

Representative immunohistochemical images of a dorsal HPC from a YinY novelty exposed animal and quantitative analysis of the effect of age, versus young, -associated GM transplantation compared to aged mice on CA3 and dentate gyrus' (DG) increase number of c-fos-positive cells after exposure to novelty (n = 3, 6, 3, 5, 4, 3 for CA3 and 4, 9, 4, 4, 4 and 7 for DG mice per group). Effect of the age-associated GM on (E) CA1 glial fibrillary acidic protein (GFAP)-astrogliosis related measures (n = 5 per group), and (F) the DG number of doublecortin (DCX) positives cells (n = 5 per group; arrows point to positive cells and the insets a representative image of a DCX positive cell in each group). (F) Representative images of the effect of human age-associated GM transplantation on CA1 increase number of c-fos-positive cells after exposure to novelty. (B-F) The color code of the different quantification experimental groups is depicted in the schematic in (A). (B-E) One-way ANOVA. Throughout the figure, bars represent the mean ± SD. N.s., non-significant; $p > 0.05$; *$p < 0.05$; **$p < 0.01$; and ***$p < 0.001$. Scale bars, 100 µm.

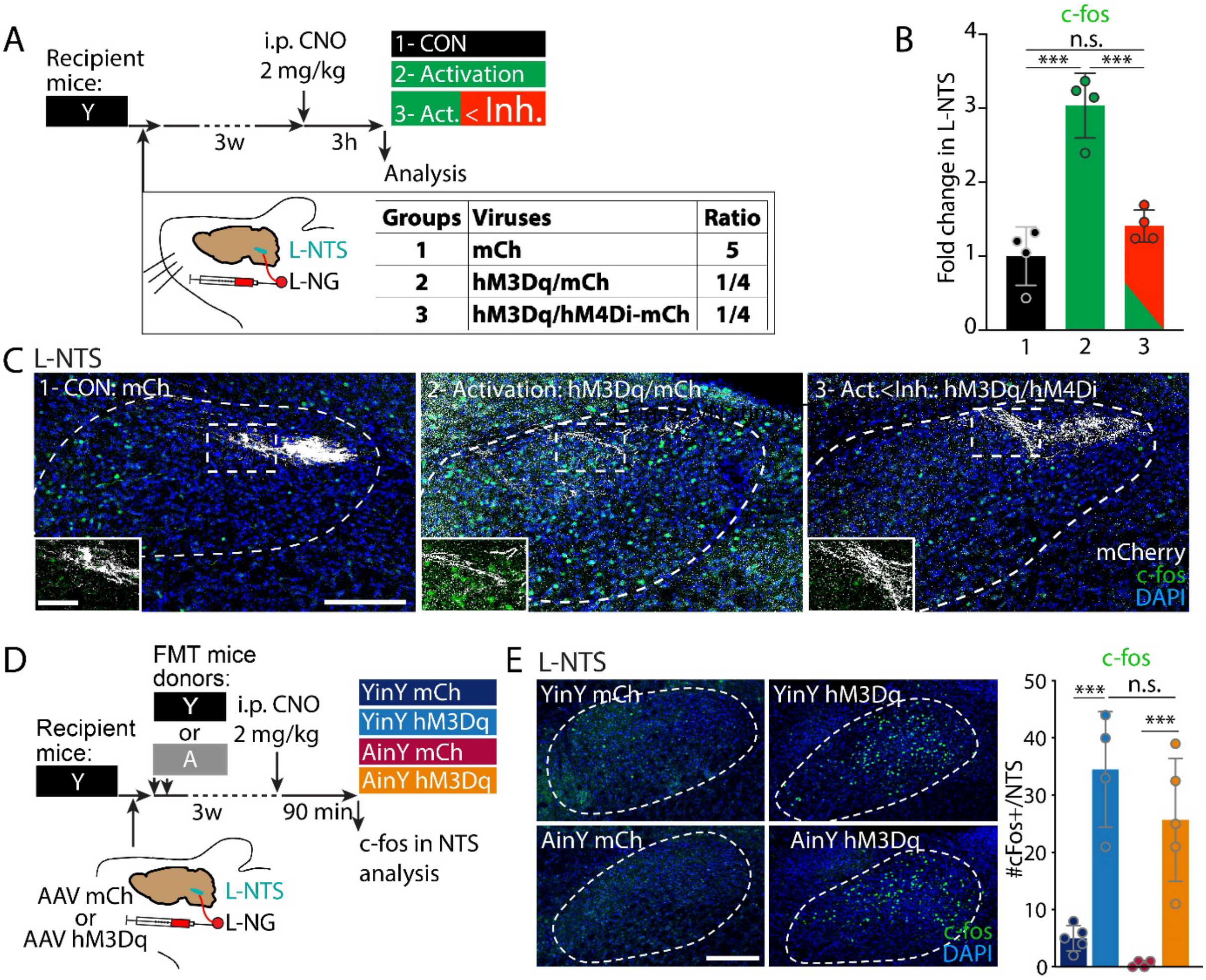


**Fig. S2. Ascending L-NG DREADD-based viral approaches efficiently promote both inhibition and activation in the ipsi-lateral NTS, and, for the latter, irrespectively of the GM-transplatation donor mice age group.** (A) Schematic of the viral and pharmacogenetic-based strategy to test the potential of hM4Di to inhibit L-NG hM3Dq ascending activation. AAV5-hSyn-DIO, -mCherry only or -hM3Dq-mCherry plus mCherry, or -hM3Dq-mCherry plus hM4Di- mCherry were co-injected (for the latter, with the inhibitory form in excess of 4 to 1 compared to the activatory one) with an AAV1-hSyn-Cre (not represented) in young mice L-NG and timing of the L-NG pharmacogenetic vagal modulation in young-adult mice. (B) quantification of the effect of L-VN hM3Dq activation and of the potential of hM4Di to counteract the VN ascending activatory effect of a DREADD activation, through quantification of the number of c-fos positive cells in the ipsi-lateral NTS of hM3Dq injected and hM4Di/hM3Dq L-NG co-injected mice (n = 4 per group) and (C) representative images of L-NG mCherry-positive ascending fibers and c-fos positive cells in the L-NTS. (D) Schematic of the viral and pharmacogenetic-based strategy to induce L-NG activation in age-associated GM transplanted mice. AAV5-hSyn-DIO-mCherry or hM3Dq-mCherry was co-injected with an AAV1-hSyn-Cre (not represented) in young mice L-NG and timing of the L-NG pharmacogenetic vagal activation in AinY mice and representative images and (E) quantification of its effect on the number of c-fos positive cells in the ipsi-lateral NTS (n = 4, 4, 4 and 5 per group). (B, E) one-way ANOVA. Throughout the figure, bars represent the mean ± SD. N.s., non-significant; ***p < 0.001. Scale bars, 100 µm, except in inset of C = 50 µm.

**Fig. S3. Graphical conclusion**

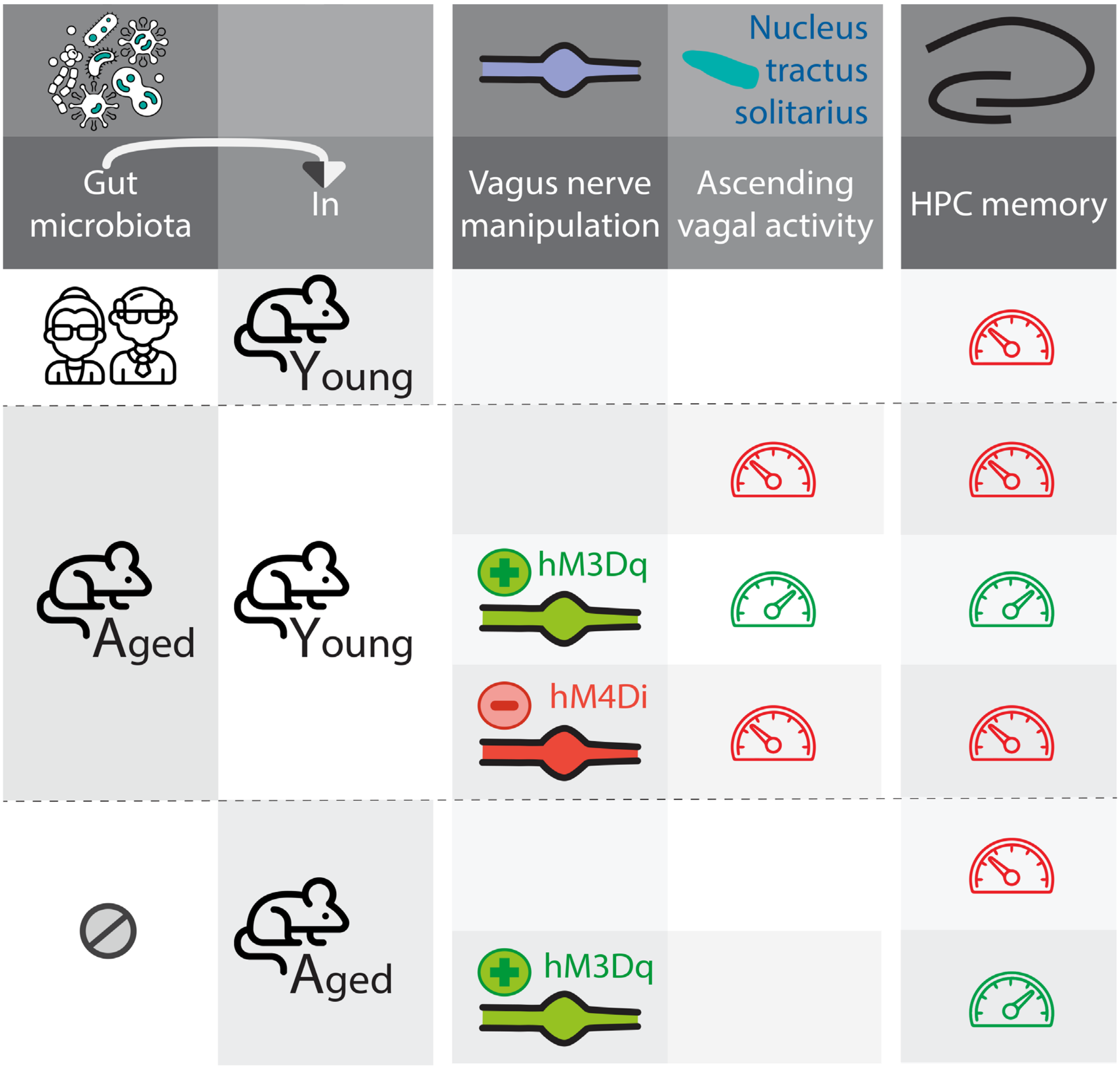

### Supplementary methods

**Fecal microbiota transplantation**. Prior to transplantation, microbiota-recipient mice received a broad-spectrum antibiotic mixture treatment of ampicillin (1 mg/ml), streptomycin (5 mg/ml) and colistin (1 mg/ml) in drinking water for 7 consecutive days combined with oral gavage with a solution of vancomycin (0,1 mg/ml) and metronidazole (0,5 mg/ml) on day 3 and 5 of the antibiotic treatment. The latter was ceased 24h before the microbiota transplantation by returning mice to regular drinking water. Fresh fecal samples (1 g in 5 mL sterile PBS) were used for the transplantation protocol, harvested from either young (Y) or aged mice (A). Recipient mice received the fecal suspension (300 mL per mouse) by oral gavage at 1- and 3-days following antibiotics discontinuation, and were maintained in isolators from one day prior to it and up to the onset of fecal matter harvest and of behavioral testing 6 and 7 weeks later, respectively.

**Effectiveness assessment of the pre-FMT antibiotic-based GM cleansing procedure.** Two fecal pellets were harvested from the 5 same mice, before and after the 7 days broad spectrum antibiotic treatment, directly in sterile 16% glycerol PBS. Samples were homogenized using 1ml plastic syringe and centrifugated for 3 min at 13.000g. Supernatant were transferred to a new tube and stored at -80°C. Samples were thawed and 5ul of this GM resuspension were mix in 100ul of sterile PBS, before being plated on a TGV-agar medium plate and incubated at 37°C for 3 days. Vehicle condition consisted in using the 16% glycerol PBS solution in place of the GM resuspension and was plated alongside the mice samples, to verify for the sterility of the solutions and procedure.

**Mice fecal harvest, DNA extraction, amplicon sequencing and analyses of 16S rRNA gene sequence data.** 6 weeks following FMT, fecal material was harvested at 9 am and stored at -80C until processing. Human donors and 6 weeks-post FMT mice fecal material was used for microbiota profiling. DNA was extracted from fecal samples using 3 mice feces and 150 mg of human material, using the RNEasy PowerMicrobiome (Qiagen) according to the manufacturer's instructions for nucleic acid extraction with the only adaptation being a 15 min vortexing for the sample lysis step. DNA concentration was normalized to 5 ng/µL by dilution with DNA elution solution (Qiagen) to produce a final volume of 20 µL. Normalized DNA samples were sent to the institute Pasteur BIOMICS center for PCR amplification of 16S RNA genes and paired-end Illumina sequencing (2 × 250 bp) on the MiSeq platform. The V4 hypervariable region of the 16S rRNA genes was amplified using the S-D-Bact-0341-b-S-17, and S-D-Bact-0785-a-A-21 primers from[43]. Following DNA extraction and sequencing, raw paired-end reads were processed in a data curation pipeline, that includes a step of removal of low-quality reads (Qiime2 2019.10)[44]. Remaining sequences were assigned to samples based on barcode matches, and barcode and primer sequences were then trimmed. Sequences were denoised using the DADA2 method, and reads were classified using Silva reference database (version 132)[45]. Alpha and beta diversity were computed using the phyloseq package (v1.24.2)[46] in R version 3.6.2. Principal Coordinate analyses of the Jaccard index were performed to assess beta diversity. The number of observed species, Chao1, Shannon and Simpson indexes were calculated using rarefied data (depth = 23,000 sequences/sample) and used to characterize alpha diversity. Raw sequence data are accessible in the European Nucleotide Archive (accession number: 2021-0023). Differential analysis was performed using the linear discriminant analysis effect size (LEfSe) pipeline[47].

**Isotropic novel-object location/recognition and fear conditioning task procedure**. Procedure was run as described [48] with minor modifications. Briefly, 7 weeks after initiation of the FMT procedure, behavior was run in the morning, mice were transferred to the testing room at least 1 h before testing. Behavioral arenas were 44 cm sides squares in which animals were habituated for 10 min per day for two consecutive days. The next day a same duration object training phase was initiated for each animal with the following objects: for ISO-NOL#1, a 31,8/15,8/9,6 mm yellow duplo brick (Lego) and a 50 ml falcon blue lid (Falcon), for ISO-NOL#2, a 35 mm diameter/40 mm high black plastic cylinder and a 40/30/20 mm aluminum rectangular cube. In both cases, the two objects were placed in the two same side of the arena-corners, 6cm away from both vertical arena walls. It was followed by a 6 min testing phase were one of the objects (counterbalanced between the different animals) was moved to the opposite, same side corner of the arena (see schematic Fig. 1A). Four arenas were used with the same number of animals ran at once. For all stages where animals were in contact with the objects, the exploratory behavior was recorded and intertrial interval was 1 hour. ISO-NOL#1 and #2 tests were run prior and 60 days following the start of the antibiotic treatment, respectively. Videos were analyzed by an experimenter blinded to the experimental treatment. Mice were scored as exploring an object if they showed obvious signs of directed attention (i.e., sniffing or contact). Time spent on top of the objects was not counted, unless the mice were simultaneously directing attention to the object. The discrimination index was calculated from by dividing the difference in the object exploration time between the familiar and new location objects with the sum of the time exploring both objects. ISO-NO and fear conditioning task was run as previously described[48].

**Novelty-exposure assay.** Animals were exposed to a novel context (an arena with a textured plastic floor) for 3 minutes, before being returned to their home cage. 90 minutes after this novelty exposure, a time corresponding to the pick of c-fos expression following a behavioral stimulus, mice were intracardially perfused with paraformaldehyde, their brains harvested and the number of c-fos positive cells in the hippocampal CA1 subregion was quantified. For the latter, automated counting was done with the ICY software (Institut Pasteur, France Bioimaging) and its 'spot detection' function in brain areas (CA1, CA3 or DG) previously drawn on the slides' images via the region of interest function. Parameters of the spot detector were set on control condition images to match the detection made manually in parallel. The same parameters were then run on all experimental conditions images and allowed an unbiased automated detection.

**Subject recruitment and human GM sample collection.** Recruitment of young and aged healthy subjects were recruited on the basis of being, penally responsible (>18 years old) at the time of inclusion, drug naïve, and either less than 35 (young adult subject group) or more than 65 (aged subjects group) years old, having an absence of systemic or familial history of medical illness or mental disorders and of antibiotics treatment less than a month before the sample collection. Sample collections were made following the standard operating procedure IHMS_SOP 003 V2 for fecal samples self-collection and laboratory analysis handled within 4 hours from the International human microbiome initiative with minor modification[49] http://www.microbiome-standards.org/index.php?id=Sop&num=003). Briefly, samples were self-collected by donors in the morning in 120 ml urinary samples collection container (Dutscher, 861086) modified to have the lid pierced to allow for gas exchange, immediately moved to an oxygen chelating plastic pouch system (Biomerieux, Genbag Anaer 45534) with the oxygen chelating pad added and sealed according to instruction of the manufacturer. The ensemble was moved to a secondary container previously stored at 4°C, containing a calorimetric cold pack to ensure a temperature control of the samples and was brought for processing less than 3 hours post-collect. Samples were aliquoted for 16S rRNA gene sequencing and for human FMT to mice, following a previously designed protocol[50], and stored at 80°C until use.

**Immunohistochemistry.** Mice were euthanized by an intra-cardiac infusion of 0.9% NaCl followed by 4% Formol under deep anesthesia (xylazine 20 mg/kg, ketamine 100 mg/kg). Brains were extracted and post-fixed overnight with Formol 4% at 4°C, received three 10-minutes PBS washes before being coronally cut into 50 µm thick sections using a vibratome (Leica), and then stored at -20°C in cryoprotective solution (27,5% sucrose, 27,5% ethylene glycol, 45% PBS). Prior to immunostaining, sections went through three 10-minutes PBS washes. The blocking buffer used to permeabilize/block and incubate the sections with the primary and secondary antibodies was a mixture of PBS with 0.2% Triton X-100 and 10% Normal Donkey Serum (Abcam, Ab7475) and Na-azide 0.01%. Immuno-labelling consisted in a blocking/permeabilization step of 1h30 under constant agitation, then primary antibodies were incubated overnight (48h for c-fos) at 4°C under constant agitation. Primary antibodies used were: c-fos (1:4000, rabbit, Sysy, 226 003; 1:2000; guinea-pig, Sysy, 226 005), GFAP (1:2000, chicken, Abcam, ab13970), DCX (1:1000, rabbit, Abcam, Ab62341), mCherry (anti-RFP, 1:4000, rabbit, Rockland, 600-401-379). After three 5-minutes washes with PBS, sections were incubated for 1h30 under agitation with either Alexa Fluor 488 or 568–conjugated secondary antibodies followed by a 5-minutes DAPI (0.001% in PBS) counterstaining wash followed by two 5-minutes PBS washes, at room temperature. Sections were then mounted on slides with a mounting medium (FluoroMount-G, Interchim). Image acquisition was carried out using a confocal microscope (LSM 700, ZEISS).

**L-NG viral transduction and pharmacogenetic modulation of ascending vagal activity.** Prior to surgery, mice received an i.p. injection of buprenorphine (0.1mg/Kg, Buprecare, Axience, Pantin, France) in saline (NaCl 0.9%), 30 minutes before anesthesia. Surgeries were performed under aseptic conditions and 1–2% isoflurane/oxygen anesthesia. Mice body temperature was maintained at 37°C using a heating pad. Surgical access to the left nodose/jugular complex was obtained by making an incision along the ventral surface of the neck and blunt dissection using surgical retractors to maintain the muscular, aortic/oesophagial neck parts and blood vessels surrounding the nodose complex region and to expose the nodose. Viral preparation used consisted of:

1) for the L-NG pharmacogenetic inhibition or activation experiments (Fig. 3 and S2E): AAV5-hSyn-DIO-hM4D(Gi)-mCherry (Addgene, titer: 4.5 X $10^{12}$ genome copies/ml) or AAV5-hSyn-DIO-hM3Dq-mCherry (Addgene, titer: 3.5 X $10^{12}$ genome copies/ml), respectively, and for control mice AAV5-hSyn-DIO-mCherry (Addgene, titer: 3.4 X $10^{13}$ genome copies/ml),
2) for the validation of the L-NG pharmacogenetic activatory and inhibitory approach (Fig. S2A-C), for group1: AAV5-hSyn-DIO-mCherry (Addgene, titer: 3.4 X $10^{13}$ genome copies/ml) only,
or a 1/4 mixture of - for group 2: AAV5-hSyn-DIO-hM3Dq-mCherry with AAV5-hSyn-DIO-mCherry (Addgene, titer: 3.5 X $10^{12}$ and 3.4 X $10^{13}$ genome copies/ml),
- for group 3: AAV5-hSyn-DIO-hM3Dq-mCherry with AAV5-hSyn-DIO-hM4D(Gi)-mCherry (Addgene, titer: 3.5 X $10^{12}$ and 4.5 X $10^{12}$ genome copies/ml).
In both cases, viral mixtures were always complemented with a 1/5 addition of AAV1-hSyn-Cre (Addgene, titer: 2.6 X $10^{13}$ genome copies/ml), and 0.05% Fast Green FCF (Sigma-Aldrich). An elongated glass capillary mounted on an Nanoject injector III (Drummond), filled with the corresponding viral mix, was inserted into the nodose/jugular complex, and 90 nl of the virus solution were injected per mouse, in three 30 nl injection bollus, at 6 nl/s. Success in injection of the nodose was determined by Fast Green Dye filling of the ganglia. Three weeks following the viral injection, pharmacogenetic ascending-vagal inhibition or activation was initiated by CNO (Enzo life sciences, Fisher scientific BML-NS105-0025) i.p. injections at 2mg/kg in saline (NaCl 0.9%), 30 min before the start of the ISO-NOL and novelty exposure procedures.
**Statistics.** All experiments and data analyses were achieved blinded. In the figures are indicated the number of subjects used in each experimental condition and the data are expressed as mean ±SD. Statistical analyses were performed with GraphPad Prism 8.0. No statistical methods were used to pre-determine sample size, or to randomize.
**Fundings.** The project leading to this application has received funding from the European Union's Horizon 2020 research and innovation program under the Marie Sklodowska-Curie grant agreement No 708466 and from the Institut Pasteur Research Applications and Industrial Relations Department grant DARRI INNOV 43-2019.
